# JAK1 inhibition blocks lethal sterile immune responses: implications for COVID-19 therapy

**DOI:** 10.1101/2020.04.07.024455

**Authors:** Kathryn D. Tuttle, Ross Minter, Katherine A. Waugh, Paula Araya, Michael Ludwig, Colin Sempeck, Keith Smith, Zdenek Andrysik, Matthew A. Burchill, Beth A.J. Tamburini, David J. Orlicky, Kelly D. Sullivan, Joaquin M. Espinosa

**Affiliations:** Linda Crnic Institute for Down Syndrome, University of Colorado Anschutz Medical Campus, Aurora, CO 80045, USA; Department of Pharmacology, University of Colorado Anschutz Medical Campus, Aurora, CO 80045, USA; Division of Gastroenterology and Hepatology, Department of Medicine, University of Colorado Anschutz Medical Campus, Aurora, CO 80045, USA; Department of Immunology and Microbiology, University of Colorado Anschutz Medical Campus, Aurora, CO 80045, USA; Department of Pathology, University of Colorado Anschutz Medical Campus, Aurora, CO 80045, USA; Section of Developmental Biology, Department of Pediatrics, University of Colorado Anschutz Medical Campus, Aurora, CO 80045, USA; Department of Molecular, Cellular and Developmental Biology, University of Colorado Boulder, CO 80305, USA

## Abstract

Cytokine storms are drivers of pathology and mortality in myriad viral infections affecting the human population. In SARS-CoV-2-infected patients, the strength of the cytokine storm has been associated with increased risk of acute respiratory distress syndrome, myocardial damage, and death. However, the therapeutic value of attenuating the cytokine storm in COVID-19 remains to be defined. Here, we report results obtained using a novel mouse model of lethal sterile anti-viral immune responses. Using a mouse model of Down syndrome (DS) with a segmental duplication of a genomic region encoding four of the six interferon receptor genes (Ifnrs), we demonstrate that these animals overexpress Ifnrs and are hypersensitive to IFN stimulation. When challenged with viral mimetics that activate Toll-like receptor signaling and IFN anti-viral responses, these animals overproduce key cytokines, show exacerbated liver pathology, rapidly lose weight, and die. Importantly, the lethal immune hypersensitivity, accompanying cytokine storm, and liver hyperinflammation are blocked by treatment with a JAK1-specific inhibitor. Therefore, these results point to JAK1 inhibition as a potential strategy for attenuating the cytokine storm and consequent organ failure during overdrive immune responses. Additionally, these results indicate that people with DS, who carry an extra copy of the IFNR gene cluster encoded on chromosome 21, should be considered at high risk during the COVID-19 pandemic.

**One Sentence Summary:** Inhibition of the JAK1 kinase prevents pathology and mortality caused by a rampant innate immune response in mice.

## Introduction

Cytokine release syndrome (CRS) or hypercytokinemia, often referred to as a ‘cytokine storm’, has been postulated to drive mortality during severe respiratory viral infections, such as influenza (*1*) - including the 1918 Spanish flu epidemic (*2*) and the H5N1 bird flu (*3*), as well as during the Severe Acute Respiratory Syndrome coronavirus epidemic of 2003 (SARS-CoV-1) (*4*). Increasing evidence supports the notion that mortality during infections with SARS-CoV-2, which causes coronavirus disease of 2019 (COVID-19), is driven by the exacerbated immune response to the virus, leading to a cascade of events involving a cytokine storm and acute respiratory distress syndrome (ARDS), often accompanied by myocardial damage and multiple organ failure (*5, 6*). This pathological cascade resembles what is observed in other lethal viral lung infections, where the immune response to the virus triggers a primary wave of cytokines, including Type I and Type III interferons (IFNs), followed by infiltration and activation of diverse immune cells and production of secondary cytokines and chemokines, including Type II IFN (IFN-*γ*), accelerated immune activation, and progressive decline of respiratory function (*7*). Importantly, many of the cytokines and chemokines involved in the cytokine storm employ Janus protein kinases (JAKs) for signal transduction.

Cytokine analysis of COVID-19 patients indicate an important role for the magnitude of the cytokine storm in prognosis and clinical outcome. Levels of C-reactive protein (CRP) and IL-6 at the time of hospitalization were reported to be significantly higher in patients who eventually died versus those that survived (*5*). In a different study, patients admitted to intensive care units (ICUs) showed significantly higher levels of IL-2, IL-10, IL-7, IP-10, MCP-1, MIP-1α, G-CSF, and TNF-α relative to non-ICU patients (*6*). Using artificial intelligence methods, high levels of circulating alanine aminotransferase (ALT), a marker of liver inflammation, was associated with progression toward ARDS (*8*). Altogether, these findings support the rationale for combining antiviral treatment and targeted immunosuppression as a therapeutic approach in COVID-19 (*9*). Accordingly, several clinical trials are currently testing the safety and efficacy of inhibitors of IL-6 signaling and JAK/STAT signaling. Furthermore, hydroxychloroquine, a molecule shown to attenuate Toll-like receptor signaling and cytokine production that is approved for the treatment of rheumatoid arthritis and systemic lupus erythematosus (*10, 11*), is also being testing in many clinical trials for COVID-19.

Within this context, we report here relevant results obtained during studies of the hyperactive immune response in a mouse model of Down syndrome (DS). We recently discovered that trisomy 21 (T21), the genetic cause of DS, causes consistent activation of the IFN transcriptional response in multiple cell types (*12-15*), which is explained by the fact that four of the six IFN receptors are encoded on chromosome 21 (*IFNAR1, IFNAR2, IFNGR2, IL10RB*). Additional investigations of the proteome (*16*), metabolome (*14*), and immune cell lineages (*13, 15*) of people with DS demonstrated that T21 causes: 1) changes in the circulating proteome indicative of chronic autoinflammation with clear dysregulation of IFN-inducible cytokines (*16*), 2) activation of the IFN-inducible kynurenine pathway, leading to elevated production of neurotoxic tryptophan catabolites (*14*), and 3) widespread hypersensitivity to IFN stimulation across the human immune system (*13, 15*). Altogether, these results support the notion that DS can be understood in good measure as an interferonopathy, whereby increased IFN signaling could account for many of the developmental and clinical impacts of T21. Now, we report results generated during our investigation of immune responses in the Dp16(1)/Yey mouse strain (referred hereto as Dp16) (*17*), a widely used mouse model of DS. The Dp16 strain carries a segmental duplication of murine chromosome 16 that is syntenic to human chromosome 21, encoding ∼119 genes, including the four Ifnrs. We found that Dp16 mice overexpress Ifnrs in all immune cell types tested, display hypersensitivity to Type I and Type II IFN stimulation, overproduce key cytokines, and display increased liver inflammation. Remarkably, Dp16 mice display lethal immune responses upon challenge with viral mimetic molecules, such as the TLR3 agonist poly-I:C [P(I:C)] or the TLR7/8 agonist imiquimod, a phenotype that is not observed in their wild type (WT) littermates or mouse strains carrying triplications of other genomic regions syntenic to chromosome 21. Strikingly, both the lethal immune phenotype and associated cytokine storm are blocked by treatment with a small molecule inhibitor of the JAK1 kinase, which also decreases liver pathology.

Overall, these results have potential far-reaching significance for the treatment of COVID-19, justifying a deeper study of JAK inhibitors as a therapeutic strategy to attenuate the cytokine storm and downstream organ failure in this condition, while also indicating that individuals with DS should be considered a high-risk population during the COVID-19 pandemic.

## Results

### Dp16 mice overexpress Ifnrs and show heightened sensitivity to IFN stimulation

In order to model the impact of T21 on chronic innate immune responses in mice, we employed the Dp16 mouse strain, which harbors triplication of a region syntenic to human chromosome 21 including four *Ifnrs* (**Figure 1A**) (*17*). To define the relationship between gene copy number and IFNR protein expression, we evaluated surface expression of IFNAR1 in peripheral immune cells from Dp16 mice by flow cytometry. Dp(10)1Yey/+ and Dp(17)Yey/+ mice (hereafter referred to as Dp10 and Dp17), which have three copies of genes that are syntenic to chromosome 21, but are disomic for *Ifnrs* (*18*), were included as controls (**Figure 1A**). Expression of IFNAR1 on the surface of CD45+ white blood cells was significantly higher in Dp16 mice, but not Dp10 or Dp17 animals, relative to WT littermates (**Figure 1B**). When different immune cell types were analyzed by flow cytometry, IFNAR1 expression was significantly higher in all cell types tested (**Figure 1C**). These results are consistent with previous reports of mRNA overexpression for all *Ifnrs* in this gene cluster in various tissues of Dp16 mice (*12, 19*).

**Figure 1.**
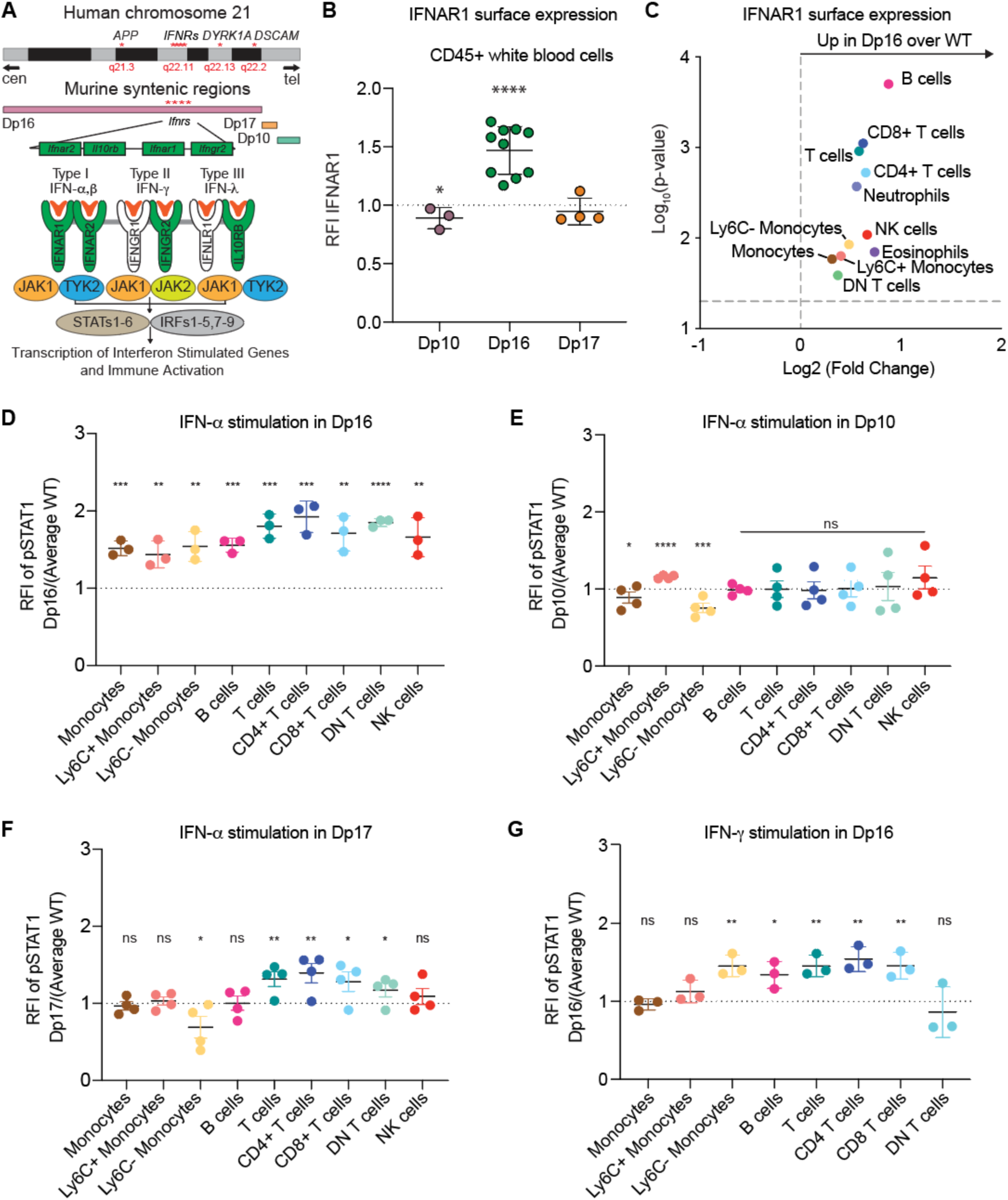
The Dp16 mouse strain is hypersensitive to interferon stimulation. **A**. Schematic of human chromosome 21, including the interferon receptor gene cluster (*IFNRs*) and syntenic regions triplicated in mouse models of Down syndrome. The *IFNR* gene cluster encodes four of the six IFN receptors: *IFNAR1, IFNAR2, IFNGR2*, and *IL10RB*. **B**. Flow cytometric analysis of IFNAR1 cell surface expression on CD45+ white blood cells (WBC) from the indicated mouse strains. Data are presented as ratios of the geometric mean fluorescence intensity (RFI) of IFNAR1 on WBC, relative to WT littermates. **C**. Volcano plot showing the fold change of IFNAR1 surface expression levels (Dp16 vs. WT) for 11 immune cell types. **D-F**. WBC from the indicated strains and WT littermates were stimulated with IFN-α (10,000 U/mL) for 30 minutes and phosphorylation of STAT1 was analyzed via flow cytometry for nine cell types. Data are presented as ratios relative to WT littermates. **G**. WBC from Dp16 and WT littermates were stimulated with IFN-γ (100 U/mL) for 30 minutes and phosphorylation of STAT1 was measured in nine cell types via flow cytometry. Data are presented as ratios relative to WT littermates. All data are presented as mean +/- SEM. Statistical significance was calculated using a Student’s T-test. *p<0.05, **p<0.01, ***p<0.001, ****p<0.0001.

To test whether upregulation of *Ifnrs* conferred stronger responses to IFN ligands, we stimulated WT and Dp16 blood samples with IFN-α and evaluated downstream phosphorylation of STAT1 (pSTAT1) by flow cytometry. Dp16 cells responded more strongly to IFN-α as defined by significantly higher levels of pSTAT1 relative to WT cells, in all immune lineages examined (**Figure 1D**). This widespread heightened response to IFN-α was not observed in Dp10 and Dp17 cells; however, T cell subsets from the Dp17 strain exhibited more robust pSTAT1 (**Figure 1E-F**). Although Dp17 mice have only two copies of the *Ifnr* genes, this result suggests that some aspect of triplication of the region of chromosome 17 syntenic to human chromosome 21 leads to differential regulation of Type I IFN signaling in T cells.

Given that one of the Type II IFN receptor subunits (*Ifngr2*) is also encoded on murine chromosome 16 and triplicated in Dp16 mice, we next investigated the response to IFN-*γ* stimulation. Indeed, Dp16 cells responded more strongly to IFN-*γ* stimulation in most cell types examined (**Figure 1G**).

Overall, these results suggest that elevated expression of the *Ifnrs* in the Dp16 mouse strain confers increased sensitivity to Type I and Type II IFNs.

### Dp16 mice display lethal immune responses to IFN-inducing TLR agonists

Since Dp16 cells mount enhanced responses to IFN ligands, we next investigated the response of Dp16 mice to innate immune stimuli known to trigger an IFN response. Chronic systemic induction of IFN signaling was elicited by repeated administration of the TLR3 agonist polyinosinic:polycytidylic acid [P(I:C)], which is known to produce an acute spike of Type I IFN production (1-3 days), followed by low but persistent expression of Type I IFN ligands and a robust cytokine response in the chronic phase (5-30 days) (*20*). WT and Dp16 mice were given intraperitoneal injections of 10 mg/kg of P(I:C) or an equivalent volume of vehicle (sham) every other day, for up to 16 days, with the experiment completed at day 17. Remarkably, Dp16 mice were profoundly sensitive to the treatment, which was largely tolerated by WT mice (**Figure 2A**). Body weight measurements during the course of the experiment revealed rapid weight loss in Dp16 mice upon treatment (**Figure 2B**). Eventually, all but one of the twelve Dp16 mice had to be removed from the study for excessive weight loss (>15%). In contrast, WT animals did not lose as much weight, and all but three out of nine survived to the end of the experiment (**Figure 2A,B**). Dp10 and Dp17 mice also tolerated chronic immune stimulation with P(I:C) (**Figure 2C-F**). Although P(I:C) treatment caused some weight loss in both Dp10 and Dp17 mice, the percentage lost was comparable to WT levels. This P(I:C)-induced weight loss was clearly dose-dependent, as mice receiving half the lethal dose (5 mg/kg) displayed only minor weight loss, with no observable differences between Dp16 and WT mice (**Figure S1**).

**Figure 2.**
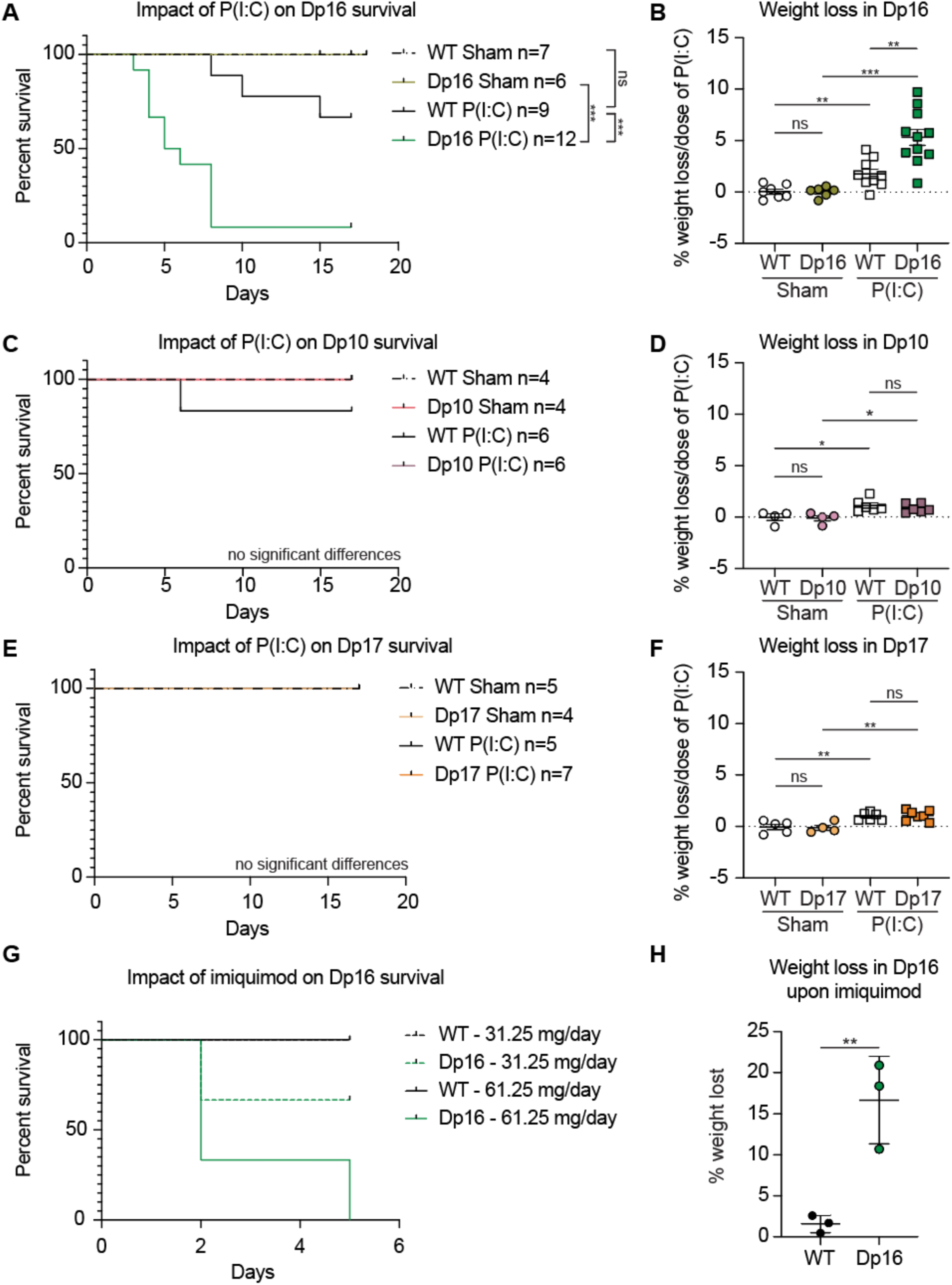
The Dp16 mouse strain displays lethal immune activation in response to viral mimetic TLR agonists. **A**. Kaplan-Meier analysis of Dp16 mice and WT littermate controls treated with 10 mg/kg P(I:C) every 48 hours for 16 days. Numbers for each group are indicated. **B**. Percentage of weight lost per injection for WT and Dp16 mice. **C** and **E**. Kaplan-Meier analysis for Dp10 and Dp17 mice, respectively, treated as in **A. D** and **F**. Weight loss analysis for Dp10 and Dp17, respectively as in **B. G**. Kaplan-Meier analysis of WT and Dp16 mice that received daily topical treatments of the indicated doses of imiquimod for up to 5 days. Data are representative of a single pilot experiment. **H**. Percentage of weight lost in WT and Dp16 mice that received 61.25 mg of imiquimod per day for up to 5 days. Data are presented as mean +/- SEM. Statistical significance for Kaplan-Meier analysis was calculated using the Mantel-Cox Log-rank test. Statistical significance for weight loss was calculated using a Student’s T-test. *p<0.05, **p<0.01, ***p<0.001, ****p<0.0001.

In order to determine whether hypersensitivity was restricted to TLR3 agonists or more broadly observed across activation of other pattern recognition receptors, we treated the mice with the TLR7 agonist imiquimod. Topical administration of imiquimod causes IFN production, acute skin inflammation, systemic inflammation, and dehydration, and is commonly employed as a model of psoriasis (*21*). Imiquimod treatment caused significant weight loss in Dp16 mice despite daily administration of supportive fluids, which led to rapid termination of the study in observation of the humane endpoint (**Figure 2G,H**). In contrast, WT animals receiving identical treatment maintained their weight (**Figure 2G,H**).

Altogether, these results indicate that the hypersensitivity to IFN stimulation observed at the cellular level in Dp16 mice associates with increased sensitivity and lethality to IFN-inducing viral mimetic agents at the organismal level.

### Dp16 mice display signs of hyperinflammation upon activation of TLR signaling

We next compared the immune response in Dp16 mice relative to their wild type littermates during the acute and chronic responses to P(I:C). First, we measured levels of circulating IFN-α 6 hours after the first injection of P(I:C) (or sham). IFN-α levels were induced by P(I:C), with a stronger response in Dp16 mice (**Figure 3A**). Elevated IFN ligand production in Dp16 mice is consistent with a predicted fast-forward loop driven by increased IFNR expression in DS (*22*). We then measured mRNA expression for several IFN-stimulated genes (ISGs) in the spleen of WT and Dp16 mice at 24 hours after a single injection of P(I:C) (or sham). This experiment revealed strong induction of ISG expression upon P(I:C) treatment, with several ISGs being more significantly elevated in the Dp16 mice than in WT animals (**Figure 3B**).

**Figure 3.**
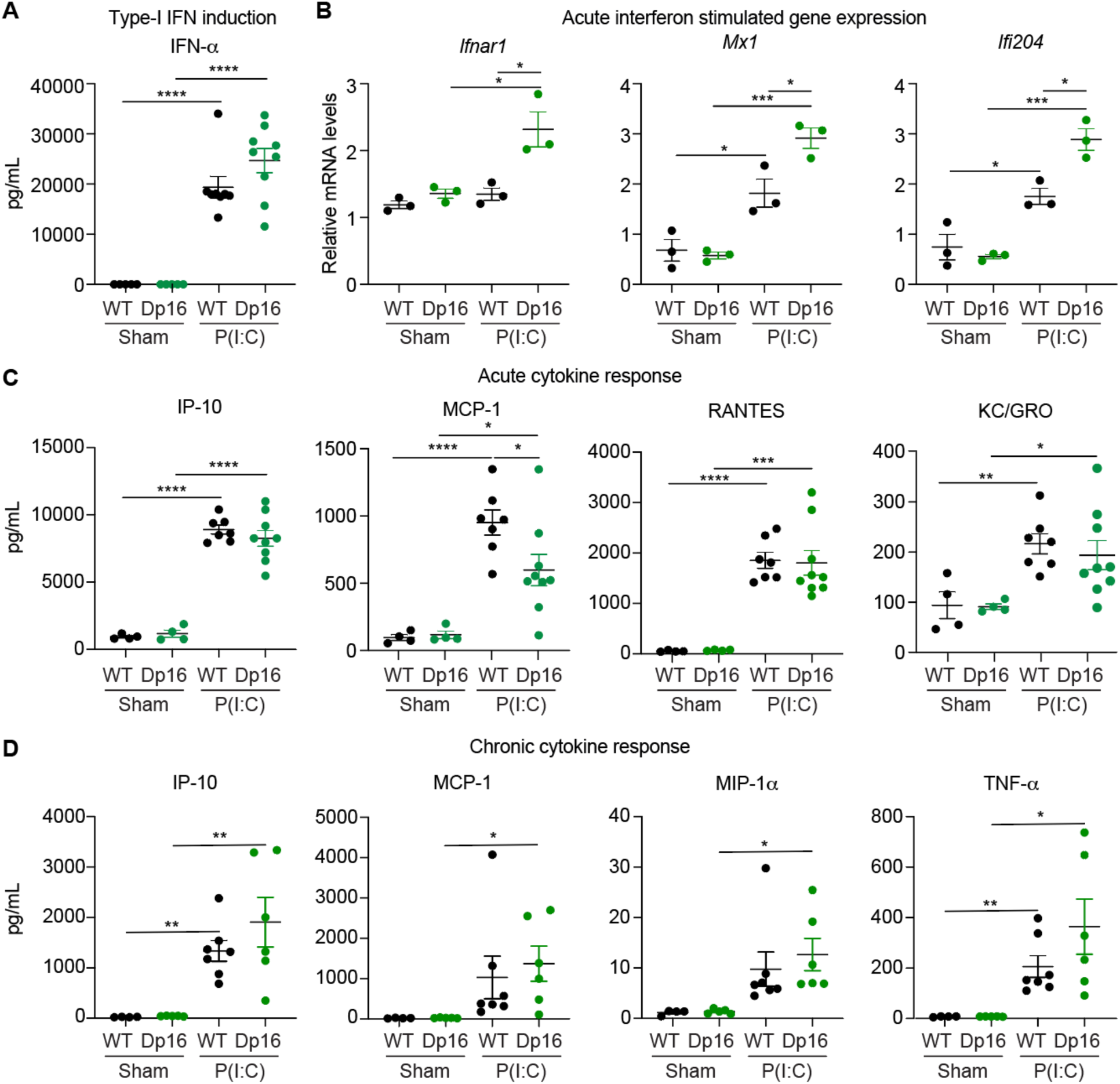
Characterization of the cytokine storm elicited by acute and chronic activation of TLR signaling. **A**. Circulating levels of IFN-α were measured 6 hours after a single P(I:C) (or sham) injection in WT versus Dp16 mice. **B**. Expression of IFN-stimulated genes (ISGs) was measured by Q-RT-PCR using RNA prepared from spleens of WT versus Dp16 mice harvested 24 hours after a single P(I:C) (or sham) injection. Expression was normalized to 18s rRNA. **C**. Circulating cytokines were evaluated 24 hours after the first P(I:C) (or sham) injection in WT versus Dp16 mice. **D**. Cytokine analysis in the peripheral blood in animals that were chronically treated with P(I:C) (or sham) as indicated. Samples from each animal were collected at either the humane endpoint of 15% weight loss or at the end of experiment at day 17, whichever occurred first. Data are presented as mean +/- SEM. Statistical significance was calculated using a Student’s T-test. *p<0.05, **p<0.01, ***p<0.001, ****p<0.0001.

Next, we measured the levels of several cytokines and chemokines in the bloodstream 24 hours after a single P(I:C) injection. Expectedly, P(I:C)-treated animals showed increased levels of many pro-inflammatory factors, regardless of genotype (**Figure 3C** and **Figure S2A**). In this acute phase, the most significantly elevated inflammatory markers were IP-10 (*Cxcl10*), MCP-1, RANTES (*Ccl5*), and KC/GRO (*Cxcl5*) (**Figure 3C**). On average, Dp16 mice did not show significant differences in expression of cytokines relative to their WT littermates. Then, we measured circulating cytokines and chemokines during the chronic response. We compared serum levels between Dp16 mice taken at the time of harvest due to reaching the humane endpoint of 15% weight loss to the levels in the WT littermates that survived to the end of the 17-day experiment. At these later stages of chronic inflammation, several of these inflammatory markers were significantly elevated in the Dp16 animals, such as MCP-1 and MIP-1α, but not in the WT littermates (**Figure 3D**).

Altogether, these results indicate that activation of TLR signaling leads to strong induction of key cytokine and chemokines in our experimental paradigm, with several inflammatory markers being more elevated in the Dp16 mice. Although not a single cytokine or chemokine stands out as the sole driver of phenotypes in the Dp16 mice, the mild over-production of several cytokines could potentially contribute to their exacerbated immune sensitivity.

### Dp16 mice show increased liver inflammation and pathology

Next, we investigated markers of liver inflammation and injury. The serum levels of the enzymes alanine transaminase (ALT) and aspartate transaminase (AST), two commonly used markers of hepatocyte injury, were significantly elevated upon P(I:C) treatment in the Dp16 mice, reaching concentrations nearly an order of magnitude higher than those observed in their WT littermates (**Figure 4A**). Prompted by these results, we investigated liver pathology. Liver tissue sections were stained with hematoxylin and eosin (H&E) and evaluated by a trained histologist, blinded against treatment group and genotype. Scoring of liver pathology was done with procedures adapted for mice (*23*) using a validated histological scoring system (*24*), that included parameters such as cell injury, inflammation, and reactive changes, leading to an overall liver pathology score. This analysis revealed that Dp16 mice have increased liver pathology even before P(I:C) treatment, as demonstrated by significantly higher levels of cell injury, inflammation, and reactive changes relative to their WT littermates (**Figure 4B,C**). This basal level of liver pathology was not observed in the Dp10 and Dp17 mice (**Figure 4D,E**). Importantly, upon treatment with P(I:C), the livers of WT animals developed pathology scores comparable or greater than those observed at baseline in Dp16 mice, including signs of strong recruitment and/or expansion of inflammatory cells. When P(I:C) was administered to Dp16 mice, all metrics of liver pathology increased, leading to significantly higher overall pathology scores relative to both Dp16 sham treatment and WT upon P(I:C) treatment (**Figure 4B-C**). Livers from Dp10 and Dp17 mice that received P(I:C) also experienced increases in liver pathology, but these changes did not differ from WT mice (**Figure 4D,E**).

**Figure 4.**
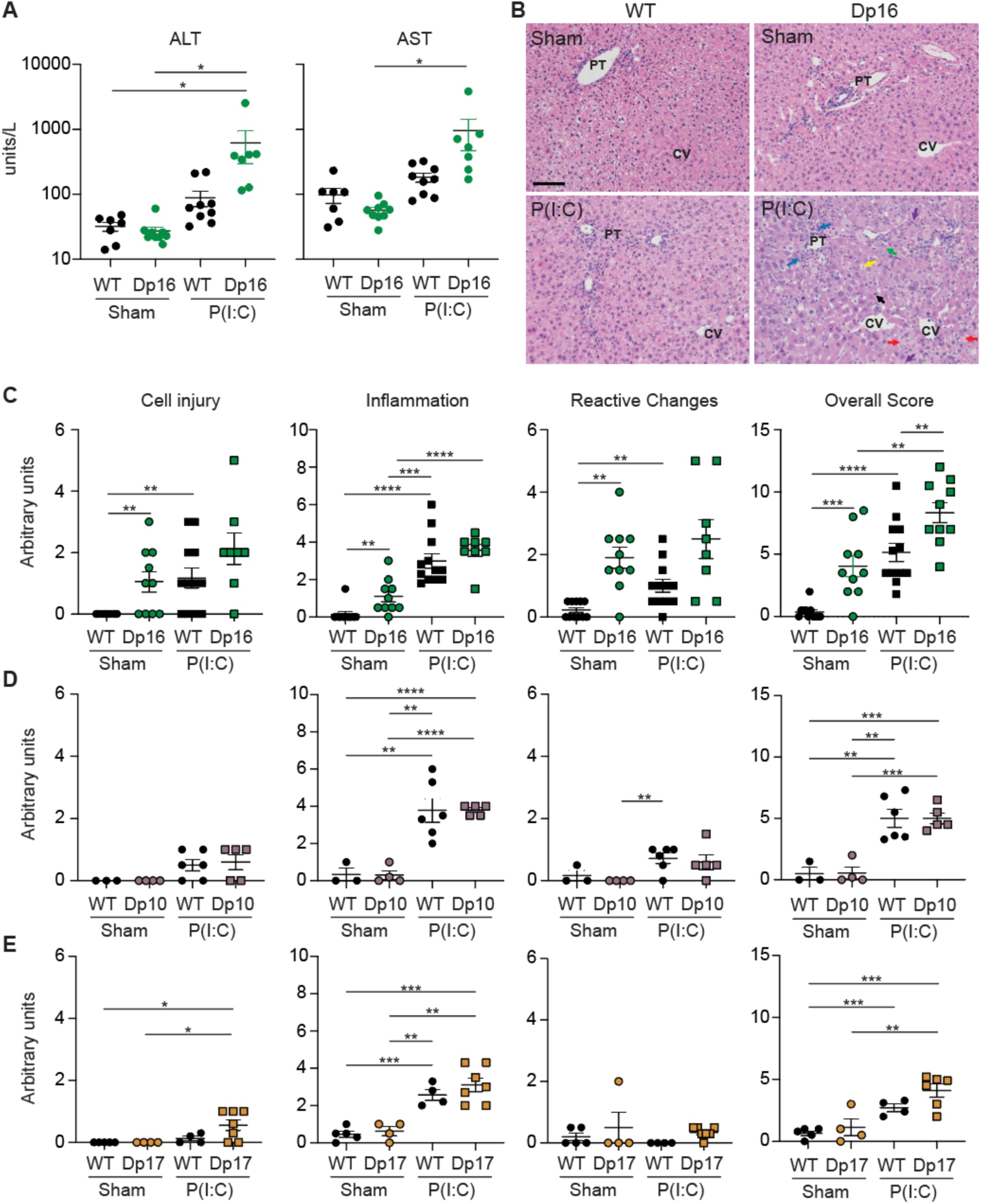
Dp16 mice display increased liver inflammation and pathology that is exacerbated by P(I:C) treatment. **A**. Comparison of alanine aminotransferase (ALT) and aspartate transaminase (AST) levels in the serum of WT and Dp16 mice, treated with vehicle (sham) or 10 mg/kg of P(I:C) for up to 16 days. Data presented as mean +/- SEM and are representative of 3 independent experiments. **B**. Representative images of H&E staining of liver sections from WT and Dp16 mice sham-treated or treated with P(I:C) as in A. The arrows indicate the following structures: red - necrotic cells, blue - inflammatory infiltrates, green - apoptotic hepatocytes, purple - circulating apoptotic cells, yellow - ballooning hepatocytes, black - mitotic bodies. PT indicates the portal triad, CV indicates the central vein. **C-E**. Metrics of liver pathology in Dp16 mice (C), Dp10 mice (D) and Dp17 mice (E) treated with vehicle (sham) or P(I:C) at the time of harvest (end of experiment at 17 days or 15% weight loss, whichever occurred first), relative to their WT littermates. Data are presented as mean +/- SEM. Statistical significance was calculated using a Student’s T-test. *p<0.05, **p<0.01, ***p<0.001, ****p<0.0001.

Overall, these results indicate that Dp16 mice display increased liver inflammation and pathology, both at baseline and upon immune activation, which could potentially contribute to their heightened sensitivity to IFN-inducing agents.

### JAK1 inhibition blocks the lethal rampant immune response

All three types of IFN signaling employ the JAK1 kinase for signal transduction, in combination with either JAK2 or TYK2 (**Figure 1A**). Therefore, we sought to determine whether inhibiting JAK1 specifically would block the lethality and pathology caused by P(I:C) treatment in Dp16 mice. To this end, we used the JAK1-specific inhibitor INCB54707. Animals received a dose of 60 mg/kg of INCB54707 (or an equivalent volume of methylcellulose used as the vehicle) twice daily via oral gavage, beginning 24 hours prior to the first P(I:C) injection, and every day during the course of the experiment. Remarkably, Dp16 mice that received the JAK1 inhibitor all survived the P(I:C) treatment, while Dp16 animals that received the vehicle and P(I:C) experienced weight loss and had to be euthanized (**Figure 5A**). We then analyzed the impact of JAK1 inhibition on cytokine production by comparing the Dp16 mice treated with P(I:C) with and without co-treatment with the JAK1 inhibitor. Importantly, cytokine production was strongly abrogated by JAK1 inhibition (**Figure 5B** and **Figure S3**). Lastly, we evaluated the impact of JAK1 inhibition on markers of liver inflammation and injury. Indeed, the JAK1 inhibitor led to a reduction of circulating levels of ALT and AST, as well as a decrease in overall liver pathology scores (**Figure 5C**).

**Figure 5.**
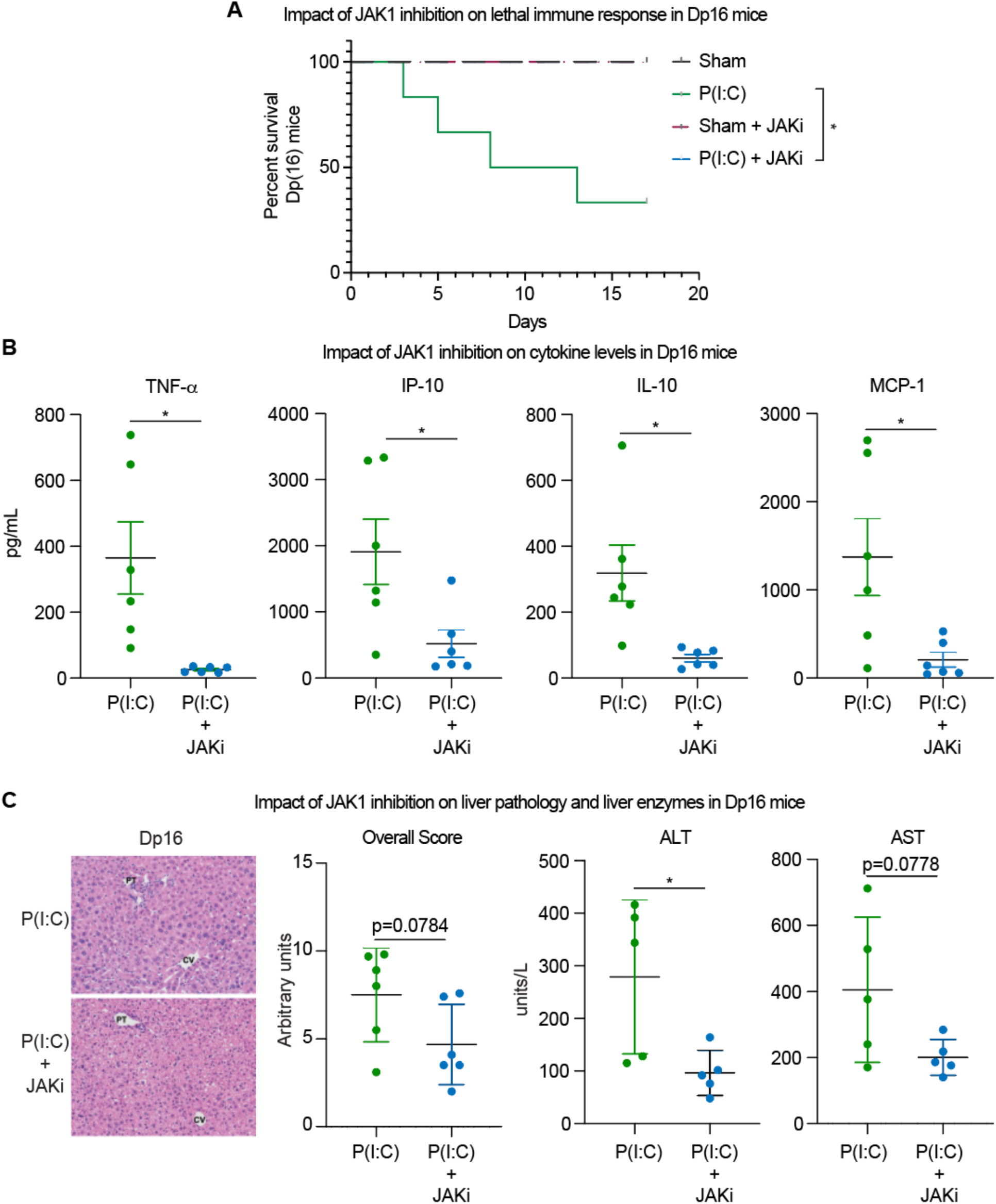
JAK1 inhibition blocks lethal immune responses, cytokine induction, and increased liver pathology in Dp16 mice. **A**. Kaplan-Meier analysis of Dp16 mice treated twice daily with INCB54707 (JAKi) or vehicle beginning 24 hours prior to treatment with P(I:C) (10 mg/kg) every 48 hours for 16 days. **B**. The indicated cytokines were measured from blood samples obtained as in Figure 3B (chronic response). P(I:C) data are same as in Figure 3B (TNF-α, IP-10, and MCP-1) and Figure S2 (IL-10), compared here to values from a parallel experimental cohort of mice co-treated with JAKi. **C**. Liver enzymes and liver pathology were assessed exactly as in Figure 4. Data are presented as mean +/- SEM. Statistical significance for Kaplan-Meier analysis was calculated using the Mantel-Cox Log-rank test and by a Student’s T-test for all other results. *p<0.05, **p<0.01, ***p<0.001, ****p<0.0001.

Altogether, these results indicate that blocking the catalytic activity of the JAK1 kinase prevents the lethality associated with a rampant innate immune response in the Dp16 mouse model. The efficacy of this treatment likely relies on a combination of inter-related effects, such as reduced cytokine production and improved liver function.

## Discussion

As the COVID-19 pandemic rapidly progresses worldwide, there is a dire need to identify new prophylactic and therapeutic strategies. Based on recent data from analyses of inflammatory markers in COVID-19 (*5, 6*) and prior knowledge about inflammatory responses in other lethal viral lung infections, targeted immune-suppression is now appreciated as a potential strategy to prevent ARDS, fulminant myocarditis, organ failure, and mortality at advanced stages of the condition (*9*). Accordingly, clinical trials for diverse immune-suppressants have been started around the world, including inhibitors of IL-6 and JAK/STAT signaling.

Here, we demonstrate a key role for the JAK1 kinase in driving cytokine storms and accompanying mortality in a mouse model of lethal anti-viral immune responses. JAK1 is one of several JAK kinases required for mediating inflammatory signaling downstream of a large number of cytokines, including all three types of IFN signaling and IL-6 (*25*). We show that eliciting the innate anti-viral immune response with TLR agonists is lethal in a sensitized mouse model carrying triplication of four of the six IFN receptors. Within days, the Dp16 mice rapidly lose weight and die if unattended, concurrent with induction of many prominent cytokines linked to the severity of COVID-19-driven pathology, such as IL-6, TNF-α, IL-10, IP-10, MCP-1, and MIP-1α (*6*). Furthermore, Dp16 mice display exacerbated liver inflammation and pathology, which, together with stronger production of some cytokines, may converge into a pathological cascade leading to wasting and death. Importantly, these phenotypes are blocked by JAK1 inhibition.

Based on these results, we propose here that JAK inhibitors are a valid therapeutic approach for attenuating the cytokine storm in COVID-19. Four JAK inhibitors are currently FDA-approved for the treatment of diverse medical conditions: Jakafi/Ruxolitinib, a JAK1/2 inhibitor approved for myelofibrosis, *polycythemia vera*, and graft-versus-host-disease; Xeljanz/Tofacitinib, a JAK1/3 inhibitor approved for rheumatoid arthritis, psoriatic arthritis and ulcerative colitis; Olumiant/Baricitinib, a JAK1/2 inhibitor approved for rheumatoid arthritis; and Rinvoq/Upadacitinib, a JAK1 inhibitor also approved for rheumatoid arthritis. Many JAK1- specific inhibitors are currently being developed and tested in clinical trials for diverse inflammatory and autoimmune conditions, including the molecule used in this study, INCB54707.

We hypothesize that JAK inhibitors could be superior agents in terms of attenuating the cytokine storm caused by COVID-19 relative to existing anti-IL6 agents, which consist of injectable monoclonal antibodies that inhibit the interaction between IL-6 and its receptor IL-6R (*26*). In contrast, JAK inhibitors are available as drugs administered orally, with very well characterized pharmacodynamics and pharmacokinetics, and may provide a more appropriate strategy to transiently tone down the cytokine storm to prevent ARDS and fulminant myocarditis. Furthermore, JAK1 inhibitors not only block IL-6 signaling, but also other inflammatory pathways involved in the cytokine storm (*25*). Depending on the degree of inhibition of various JAK kinases, it is predicted that available JAK inhibitors will block various aspects of the cytokine storm, with potential for varying degrees of therapeutic benefit.

With regards to its relevance to therapeutic strategies for COVID-19 and other conditions associated with cytokine storms, this study has several limitations. First, our mouse model does not fully reproduce the human conditions associated with cytokine storms. Although intraperitoneal injections of P(I:C) induce a clear cytokine storm in mice, this may be different in quality and magnitude to what is observed in humans. Second, there is no active viral infection or adaptive immune responses in our model. Furthermore, the genomes of SARS-CoV-2 and other lethal viruses that induce cytokine storms encode for proteins that modulate the immune response, including anti-IFN factors (*27, 28*), and it is unclear how these viral factors may modulate the course of disease and response to JAK inhibitors. Lastly, there may be significant differences in the course of a cytokine storm initiated from the lung tissue, as opposed to the more systemic response elicited in our model. Nevertheless, our results justify a thorough analysis of JAK inhibitors in pre-clinical models and clinical trials in the ongoing efforts to attenuate the impact of the COVID-19 pandemic.

Additionally, our results indicate that people with DS, who carry an extra copy of the four IFNRs encoded on chromosome 21, should be considered a high-risk population during the COVID-19 pandemic. Our studies of the Dp16 mice were originally driven by our interest in studying immune dysregulation and IFN hyperactivity in DS. Of note, our results are consistent with studies of SARS-CoV-1 demonstrating that Type I IFN signaling contributes to SARS-driven pathology and mortality in mice (*29*). We predict that, as has been observed during RSV (*30*) and H1N1 (*31*) infections, people with DS will develop more severe complications upon SARS-CoV-2 infection, including higher rates of hospitalization and mortality. We hope these results will encourage special attention to individuals with DS, including closer monitoring of hyperinflammation during COVID-19, and inclusion in clinical trials for targeted immune-suppressants.

## Materials and Methods

### Study design

This study originated from our interest in understanding immune responses and IFN hyperactivity in mouse models of DS. All animal experiments were designed *a priori*, including proper calculation of numbers of mice based on power analyses, and reviewed and approved by the Institutional Animal Care and Use Committee (IACUC) at the University of Colorado Anschutz Medical Campus. All molecular and cellular biology experiments were performed with validated reagents by trained personnel. All data generated is made publicly available in the enclosed Supplemental Materials.

### Mice

The Dp(10Prmt2-Pdxk)1Yey/J (Dp(10)1Yey/ +), Dp(16Lipi-Zbtb21)1Yey/J (Dp(16)1Yey/ +), and Dp(17Abcg1-Rrp1b)1(Yey)/J (Dp(17)1Yey/ +) strains have been previously described (*18*). Dp16 mice were purchased from Jackson Laboratories or provided by Drs. Faycal Guedj and Diana Bianchi at the Eunice Kennedy Shriver National Institute of Child Health and Human Development. Animals were maintained on the C57BL/6J background and housed in specific pathogen-free conditions.

### Chronic P(I:C) treatment

Vaccigrade P(I:C) (high molecular weight) was purchased from Invivogen and reconstituted in saline according to the manufacturer’s instructions. Mice were injected with P(I:C) at 10 mg/kg of body weight at 2-day intervals for up to 16 days. Weight was monitored throughout the duration of the experiment. Animals were sacrificed one day after the final dose (day 17) or when they lost more than 15% of their body weight. For molecular analysis, tissues were minced into small pieces and homogenized for 20 seconds in matrix Lyse-D tubes containing RLT and 0.143M *β*-mercaptoethanol.

### INCB54707 (JAKi) treatment

Mice were treated with 60 mg/kg of INCB54707 (or an equivalent volume of 0.5% methylcellulose vehicle) twice daily via oral gavage, beginning 24 hours prior to the first P(I:C) injection, and every day during the course of the experiment.

### IFN stimulations and intracellular staining

Peripheral blood was collected from the submandibular vein into Lithium Heparin tubes. 25 μL of blood were transferred to a 96-well plate, briefly subjected to RBC lysis, and then re-suspended in 30 μL of media containing 10,000 units/mL of recombinant IFN-α2a or 100 units/mL of recombinant IFN-*γ* (R&D Systems). Antibodies against SiglecF, Ly6C, CD115 and CD11b and mouse FC receptors were also included in the stimulation media. Cells were stimulated for 30 minutes at 37°C, washed once in cold FACS buffer, then fixed in 200 μL of BD Lyse-Fix buffer for 10 minutes at 37° C. Cells were permeabilized for 30 minutes in ice-cold permeabilization buffer III (BD phospho-flow).

Following permeabilization cells were stained with the following antibodies conjugated to fluorophores: CD3 (17A2), CD4, CD8 (53.6.7), B220, Ly6C, MHCII, NK1.1 and phospho-STAT1 (Tyr701).

### Flow cytometry

Whole blood was lysed gently in RBC lysis buffer, then stained for surface markers in FACS buffer at 4°C for 30 minutes. Cells were washed twice in FACS buffer, fixed in 4% PFA for 10 minutes. Fixative was washed out and cells were analyzed by flow cytometry using either a BD LRS-Fortessa X-20 or the Cytek Aurora spectral cytometers. Antibody information is provided in **Table S1**.

### Q-RT-PCR

Q-RT-PCR was performed using the Applied Biosystems Viia7™ 384-well block real time PCR system. Q-RT-PCR master mix was prepared with Applied Biosystems SYBR Select Master Mix for CFX. Standard curves were run for every primer pair in each Q-RT-PCR experiment to ensure efficient amplification of target transcripts within all experimental tissues. All samples were run in triplicate, averaged and normalized to 18s rRNA. Primer sequences at provided in **Table S2**.

### Cytokine measurements

Blood cytokine levels were measured using Mesoscale Discovery Assays and/or Biolegend Legend plex assays per manufacturer’s instructions. All samples were analyzed in duplicate and the average used for statistical analysis. Missing values were set to the lower limit of detection.

### Liver Histopathology

Formalin-fixed paraffin-embedded pieces of liver were sectioned at 5 microns and stained with hematoxylin and eosin (H&E). Scoring of liver pathology used procedures adapted for mice as described (*23*) from the validated histological scoring system established by Kleiner and Brunt (*24*).

### Statistical analysis

All statistical analysis was done with the Prism software and the exact tests employed are described in the legends for each figure. Statistical significance for Kaplan-Meier analysis was calculated using the Mantel-Cox Log-rank test. All other p values were calculated by Student’s T-test for all other results. In all cases, statistical significance is indicated as follows: *p<0.05, **p<0.01, ***p<0.001, ****p<0.0001.

## Supplementary Materials

**Figure S1.**
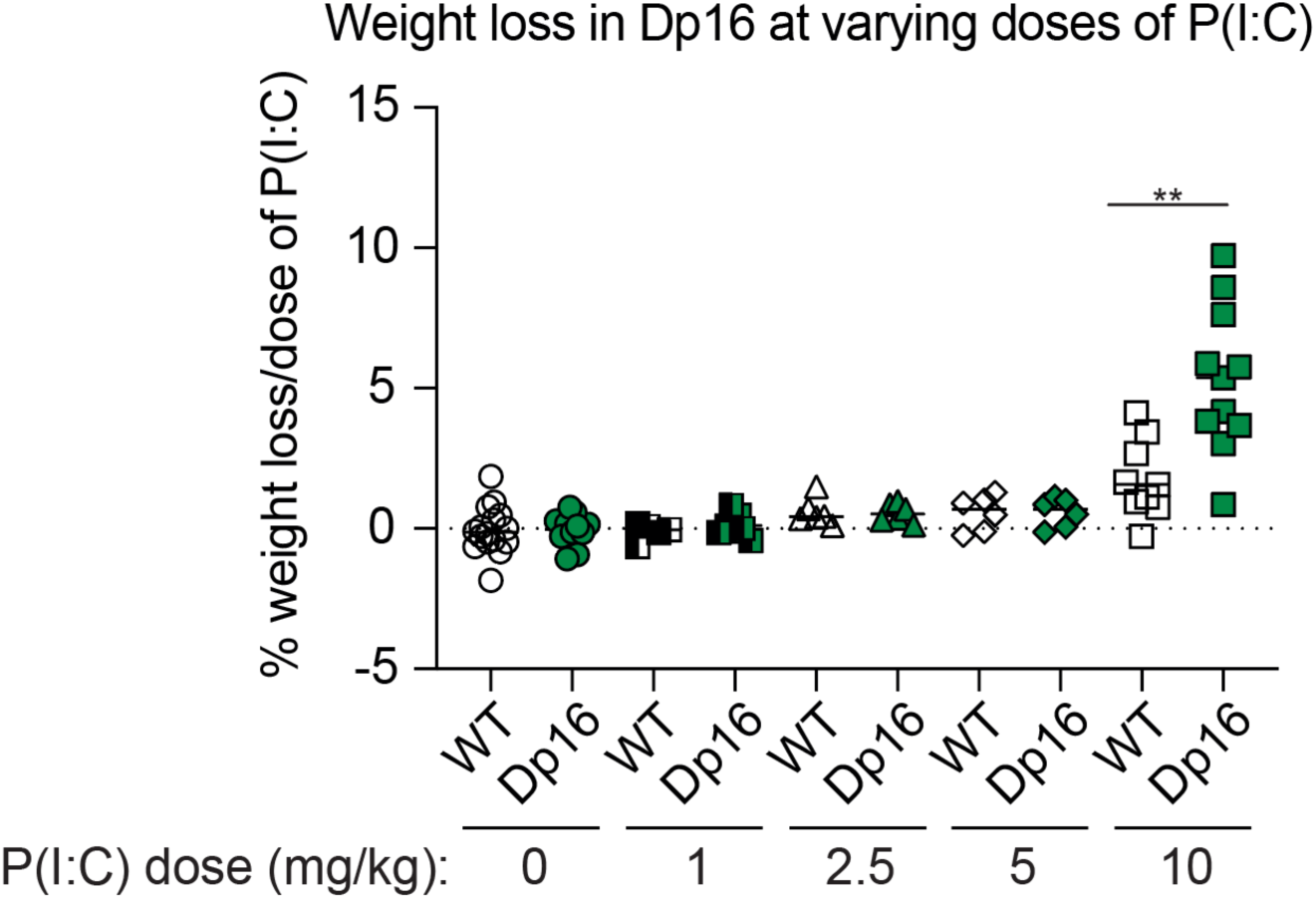
Percentage of weight lost per injection of P(I:C) in WT and Dp16 mice at the indicated doses of P(I:C). Data are presented as the mean +/- SEM and are representative of at least two independent experiments. Statistical significance for Kaplan-Meier analysis was calculated using the Mantel-Cox Log-rank test. Statistical significance for weight loss was calculated using a Student’s T-test. **p<0.01

**Figure S2.**
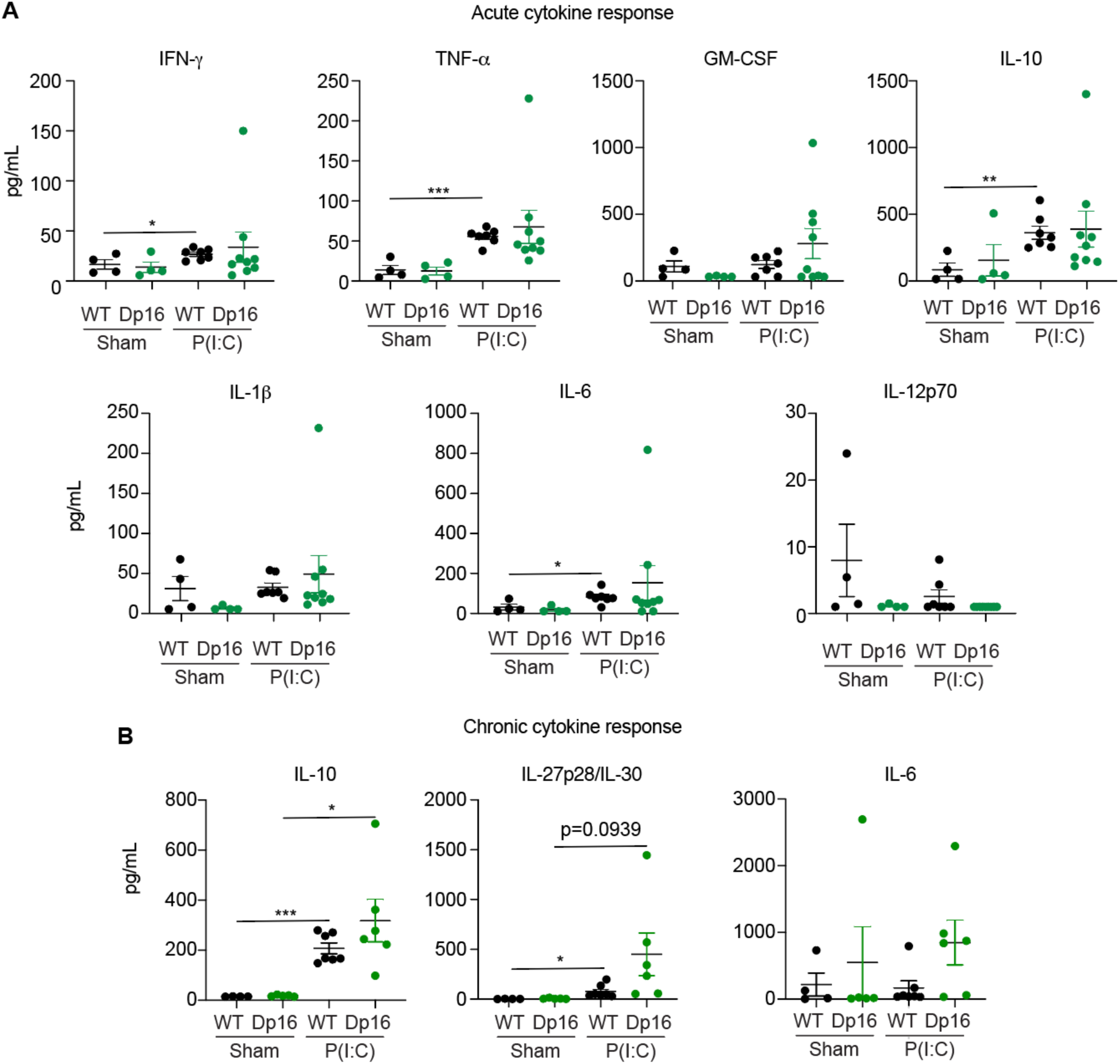
Induction of cytokines and chemokines in Dp16 mice and WT littermates. **A**. Blood samples were obtained 24 hours after the first P(I:C) injection (or sham) and evaluated for the indicated cytokines in WT versus Dp16 mice. **B**. Blood samples were collected at either at the humane endpoint of 15% weight loss or at the end of experiment at day 17, whichever occurred first, and evaluated for the indicated cytokines in WT versus Dp16 mice. Data are presented as mean +/- SEM. Statistical significance was calculated using a Student’s T-test. *p<0.05, **p<0.01, ***p<0.001, ****p<0.0001.

**Figure S3.**
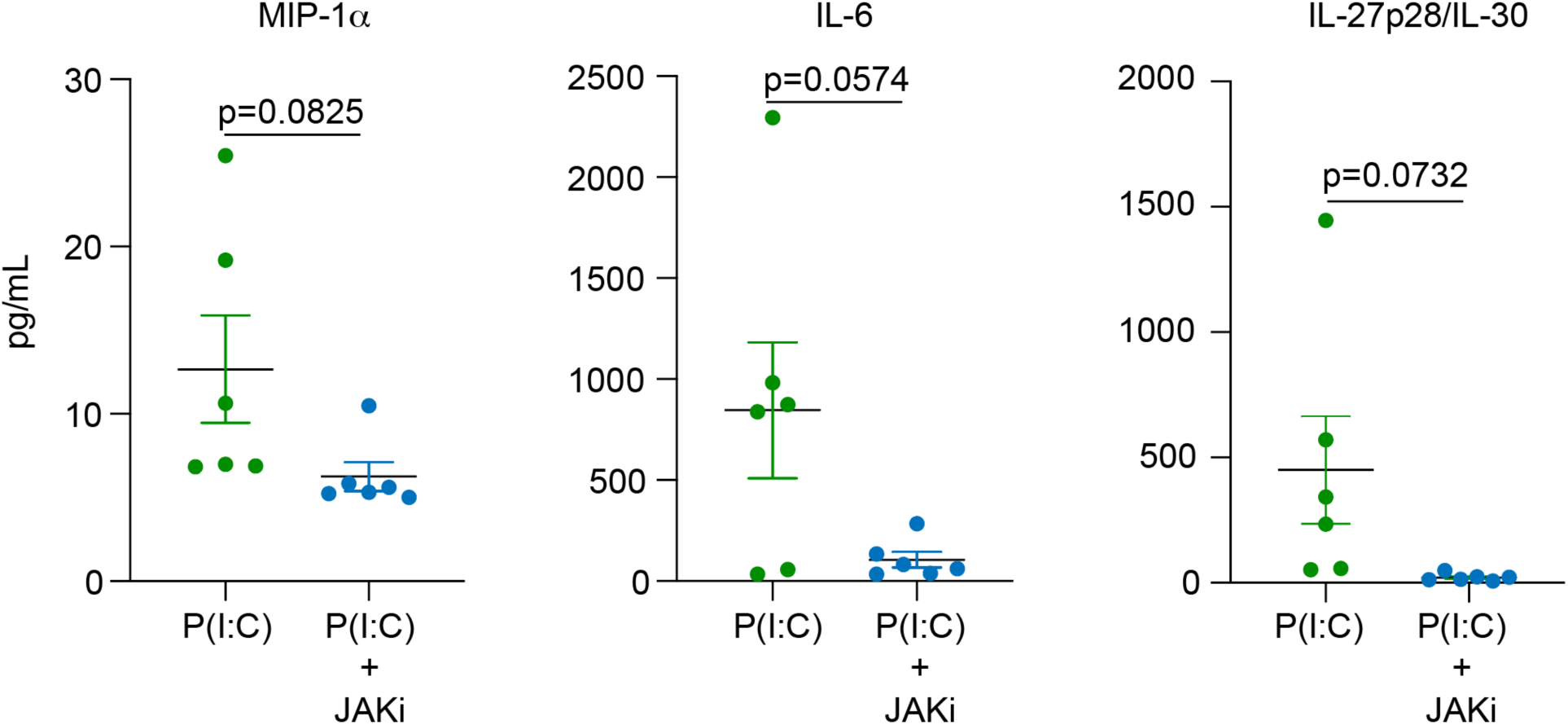
JAK1 inhibition blocks the cytokine storm in Dp16 mice. The indicated cytokines were measured from blood samples obtained as in Figure 3B (chronic response). P(I:C) data are same as in Figure 3D (MIP-1α) and Figure S2 (IL-6 and IL-27p28/IL-30), compared here to values from a parallel experimental cohort of mice co-treated with INCB54707 (JAKi). Data are presented as mean +/- SEM. p values were calculated by a Student’s T-test.

**Table S1.** Table of Antibodies. Table showing antibodies used, fluorophores, clone number, manufacturers, and catalog number.

| **Antibody target** | **Fluorochrome** | **Clone** | **Catalog #** | **Manufacturer** |
| --- | --- | --- | --- | --- |
| I-A/I-E | BUV737 | M5/114 | 748845 | BD Biosciences |
| CD11b | BUV395 | M1/70 | 563553 | BD Biosciences |
| CD4 | BV711 | RM4-5 | 100557 | Biolegend |
| Ly6C | BV605 | HK1.4 | 128036 | Biolegend |
| CD115 | BV421 | AFS98 | 135513 | Biolegend |
| CD45 | Pacific Blue | 104 | 109820 | Biolegend |
| IFNAR1 | PE | MAR1-5A3 | 127312 | Biolegend |
| STAT1 pY701 | PE | Clone 4a | 612564 | BD Biosciences |
| CD11b | Alexa Fluor 532 | M1/70 | 58-0112-82 | Life Technologies |
| NK1.1 | BB700 | PK136 | 566502 | BD Biosciences |
| CD3 | PeCy7 | 145-1C11 | 100319 | Biolegend |
| CD3 | PeCy7 | 17A2 | 100219 | Biolegend |
| CD8 | APC | 53-6.7 | 100712 | Biolegend |
| CD8 | BV510 | 53-6.7 | 100751 | Biolegend |
| Ly6G | APC-Cy7 | 1A8 | 127623 | Biolegend |
| Ly6G | APC-Cy7 | 1A8 | 25-1276-U025 | Tonbo |
| B220 | Alexa Fluor 700 | RAE3-6B2 | 103254 | Biolegend |
| Siglec F | BB515 | E50-2440 | 566211 | BD Biosciences |
| I-A/I-E | BV650 | M5/114 | 107641 | Biolegend |
| CD8 | BUV395 | 53-6.7 | 565968 | Biolegend |
| Viability | Efluor506 |  | 65-0866-14 | eBioscience |
| CD44 | BV650 | IM7 | 103049 | Biolegend |
| CD8 | BUV737 | 53-6.7 | 612759 | BD Biosciences |
| I-A/I-E | Alexa Fluor 700 | M5/114 | 107621 | Biolegend |
| CD16/CD32 | Unconjugated | 93 | 14-0161-82 | eBioscience |

**Table S2.**
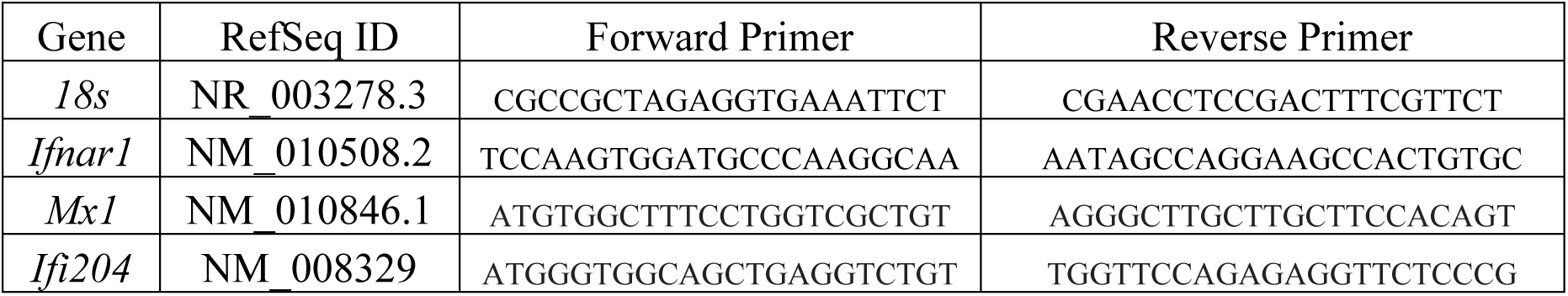
Primers for Q-RT-PCR. Primers used for Q-RT-PCR analysis of interferon-stimulated genes and 18s rRNA, including Gene, RefSeq ID, forward primer sequence, reverse primer sequence.

## Acknowledgments

We are grateful to all staff at the Linda Crnic Institute for Down Syndrome, specially Hannah Dougherty, Jessica Baxter, Dayna Tracy, and Ross Granrath. We also thank staff at the University of Colorado Cancer Center Flow Cytometry Shared Resource, the Human Immune Monitoring Shared Resource (HIMSR), and the Barbara Davis Center Flow Cytometry Core who assisted with various aspects of this work. We are grateful to Incyte Corporation for providing INCB54707 for these studies and for expedited review of these results prior to publication.

## Author contributions

KT, RM, KW, PA, ML, CS, KS, and ZA designed experiments and analyzed data. MAB, BAJT, and DJO characterized, documented and interpreted results on liver inflammation and pathology. KDS and JME designed experiments, analyzed data and provided oversight and intellectual leadership to the project. KT, KDS and JME wrote the manuscript. All authors reviewed the manuscript and made important edits and suggestions.

## Data and materials availability

All data used to generate the figures will be provided in a supplementary Data File and deposited in public repository.

